# DEFECTIVE KERNEL1 (DEK1) regulates cellulose synthesis and affects primary cell wall mechanics

**DOI:** 10.1101/2022.11.29.518391

**Authors:** Lazar Novaković, Gleb E. Yakubov, Yingxuan Ma, Antony Bacic, Kerstin G. Blank, Arun Sampathkumar, Kim L. Johnson

**Author notes:** School of Biology, Faculty of Biological Sciences, University of Leeds, Woodhouse Lane, LS2 9JT, Leeds, United Kingdom (Arun Sampathkumar, Kim L. Johnson). Corresponding authors, Kim Johnson, AgriBio building, La Trobe University, 5 Ring road, Bundoora, Victoria, Australia, 3086.

## Abstract

The cell wall is one of the defining features of plants, controlling cell shape, regulating growth dynamics and hydraulic conductivity, as well as mediating plants interactions with both the external and internal environments. Here we report that a putative mechanosensitive Cys-protease DEFECTIVE KERNEL1 (DEK1) interacts with cell wall integrity (CWI) pathways and regulation of cellulose synthesis. Our results indicate that DEK1 is an important regulator of cellulose synthesis in epidermal tissue of *Arabidopsis thaliana* cotyledons during early post-embryonic development. DEK1 is involved in regulation of cellulose synthase complexes (CSCs) by modifying their biosynthetic properties, possibly through interactions with various cellulose synthase regulatory proteins. Mechanical properties of the primary cell wall are altered in *DEK1* modulated lines supporting a role in maintenance of CWI. DEK1 affects stiffness of the cell wall and thickness of the cellulose microfibrils bundles in epidermal cell walls of cotyledons.

## INTRODUCTION

The polysaccharide rich extracellular matrix of plant cells have crucial roles in providing mechanical support, enabling water transport, cell-to-cell communication, and are a determining factor for turgor driven morphogenesis (Cosgrove, 2005). A significant component of most types of cell walls is cellulose, organized into cellulose microfibrils that impart mechanical strength to the cell wall (CMF) (Johnson et al., 2018; Zhang et al., 2019). Cellulose is synthesized by membrane bound CELLULOSE SYNTHASE (CESA) complexes (CSCs) organized in heterotrimers (Persson, 2007; Nixon et al., 2016; Polko and Kieber, 2019). CESAs are transported to the plasma membrane (PM) in vesicles originating from the Golgi complex (Sampathkumar et al., 2013) and inserted into the PM by exocytosis, mediated by a large complex of proteins called the exocyst. Once inserted into the PM, CESAs associate with cortical microtubules which is mediated by CELLULOSE SYNTHASE INTERACTING1 (CSI1/POM2) proteins (Bringmann et al., 2012). Microtubules serve as guide tracks for the CSCs. Cellulose chains synthesized by CESAs coalesce into CMF in the apoplast as they are extruded into the amorphous matrix of the cell wall, which in primary walls of dicots largely consists of two other major classes of polysaccharides, pectins and hemicelluloses (Johnson et al., 2018). Once extruded, CMF get entangled into the cell wall matrix and their continuous assembly by the CESAs moves the CSCs backwards in the PM (Diotallevi and Mulder, 2007). Mutants for either different *CESA* genes, such as *prc1-1*, or for regulatory proteins, such as *csi1/pom2* and *korrigan* (*kor*), have decreased CSC motility resulting in decreased cellulose content (Fagard et al., 2000; Lane et al., 2001; Bringmann et al., 2012; Vain et al., 2014). In addition, post-translational modifications of CESAs, such as phosphorylation, are also found to affect CSC mobility (Sánchez-Rodríguez et al., 2017).

Changes in cell wall composition and interaction between its component polymers greatly affect its mechanical properties and in turn impact growth and development of the plant (Zhang et al., 2019; Wang et al., 2020). Mutants for pectin synthesis, such as either *quasimodo2* or *xyloglucan xylosyltransferase 1, 2* (*xxt1xxt2*) have less stiff cell walls than wild type (WT) plants. Lack of either pectin or xyloglucans (XGs) in these mutants decreases bundling of CMFs, changing the mechanical properties of cell walls and initiating complex signaling cascades that result in developmental defects (Xiao et al., 2016; Du et al., 2020). Decreased synthesis of pectins and XGs also results in slower CSCs and less cellulose being produced, demonstrating that all components of the cell wall contribute to maintenance of its biochemical composition and integrity, in a continuous feedback loop-like manner (Xiao and Anderson, 2016; Du et al., 2020).

Proteins capable of sensing changes in biomechanical properties/mechanical signals at the level of the apoplast (PM/cell wall) are important for maintenance of cell wall integrity (CWI) (Engelsdorf and Hamann, 2014; Vaahtera et al., 2019). Previous studies have identified several protein families involved in CWI sensing and monitoring mechanical status of the cell wall. These proteins belong to a family of *Catharanthus roseus* receptor like kinases (*Cr*RLK), wall associated kinases (WAKs) and mechanosensitive (MS) ion channels (Vaahtera et al., 2019). Some of these proteins (eg. WAK2 or FERONIA (FER)) interact directly with pectins via their extracellular domains, while some, such as MS ion channels, sense mechanical changes in the PM caused by mechanical perturbations of the cell wall. Changes in cell wall mechanics caused by either different developmental processes or environmental stress factors are sensed by these CWI sensors, eliciting different intra- and extra-cellular responses (Kohorn, 2009; Feng et al., 2018; Gjetting et al., 2020). Rapid mechanically activated (RMA) calcium currents are known to be activated by mechanical signalling. A functioning DEFECTIVE KERNEL1 (DEK1) protein has been shown to be crucial for RMA activity in *Arabidopsis* (Tran et al., 2017). DEK1 has been identified as a regulator of cell wall biochemical composition and structure (Johnson, 2008; Demko et al., 2014; Amanda et al., 2016; Amanda et al., 2017; Galletti et al., 2015). Roles in both cell wall regulation and transduction of mechanical signals points to a possible role of DEK1 in CWI sensing pathways.

DEK1 belongs to the calpain superfamily of regulatory proteases. Calpains function in a range of cellular signaling pathways by modulating their targets through proteolytic processing resulting in altered protein activity, localization, substrate specificity or stability (Ono and Sorimachi, 2012). DEK1 is the only identified member of the calpain family in land plants (Lid et al., 2002; Demko et al., 2014) and its overall structure is distinct from the mostly cytosolic animal calpains (Lid et al., 2002; Wang et al., 2003). DEK1 is an approx. 240 kDa transmembrane protein with a complex structure including 21-23 transmembrane domains (TM), a LOOP domain, a cytoplasmic regulatory JUXTAMEMBRANE (JUXTA) domain and a proteolytically active CALPAIN domain close to its C-terminus (Lid et al., 2002; Johnson, 2008). Complementing *dek1* mutant lines of either *Arabidopsis* or *Physcomitrella patens* with only the CALPAIN domain is sufficient to rescue mutants, confirming the CALPAIN region as the catalytically active domain of the protein (Johnson, 2008; Demko et al., 2014). The catalytic activity of DEK1 CALPAIN domain is dependent on calcium (Wang et al., 2003), as shown for animal calpains (García Díaz et al., 2006). Upon cleavage, CALPAIN is released into the cytoplasm where it interacts with its yet unidentified targets (Amanda et al., 2016; Johnson et al., 2008).

DEK1 is a major regulator of plant development and growth (Johnson et al., 2005; Johnson et al., 2008). Loss-of-function mutants of *dek1* result in embryo lethal phenotypes in both *Arabidopsis* and maize (Becraft et al., 2002; Johnson et al., 2005). Different *dek1* RNA interference, artificial microRNA (amiRNA) lines with reduced levels of DEK1 or lines overexpressing CALPAIN domain have shown a range of severe phenotypes, such as an almost complete absence and misspecification of the epidermal layer (Johnson et al., 2005), an irregular epidermal layer made of large cells (Johnson et al., 2008) or a loss of cell-to-cell adhesion and occasional gaps in epidermal layer (Galletti et al., 2015). In addition, DEK1 indirectly controls HD-ZIP IV transcription factors which are involved in the regulation of epidermal identity (Galletti et al., 2015). It was proposed that DEK1 could act as a mediator of cellular adhesion by receiving and integrating signals from the apoplast and facilitating cell-to-cell contact through maintenance of epidermal identity (Galletti et al., 2015)

Studies performed on an overexpressor for *CALPAIN* (*OE CALPAIN*) and on *amiRNA DEK1* lines have shown that DEK1 is a major regulator of cell wall composition and structure in the epidermis of *Arabidopsis* leaves (Amanda et al., 2016). Immunohistochemical studies on *OE CALPAIN* lines showed a significant increase in antibody density for both cellulose, high- and low-methyl esterified homogalacturonan (HG) and arabinan while the same cell wall components were less abundant in epidermal cell walls of *amiRNA DEK1* compared with the WT (Amanda et al., 2016).

Increased labelling of epitopes for crystalline cellulose were specifically observed in epidermal cell walls of leaves and stems with no change in overall cellulose content in leaves of *OE CALPAIN-GFP* in a *dek1-3* mutant background and an *amiDEK1* line (Amanda et al., 2016). In this study, we investigated the role of DEK1 in the regulation of cellulose synthesis at a molecular level and possible roles in CWI sensing. An overexpressor of *CALPAIN* domain of *DEK1* in WT background (*pRPS5A:CALPAIN-6XHIS, OE CALPAIN*, Galletti et al., 2015) and an EMS single point mutant, *dek1-4*, which has a C to T nucleotide substitution in the CALPAIN domain and has mild phenotypes (Roeder et al., 2012) were used in this study to enable more direct comparisons to WT plants. Results of this study demonstrate that DEK1 is involved in cellulose synthesis in *Arabidopsis* cotyledon pavement cells by regulating the dynamics of CSCs. Our data also indicate that DEK1 is influencing cell wall mechanics, as well as participating in CWI sensing.

## RESULTS

### DEK1 modulated lines show altered responses to Isoxaben-induced cellulose perturbations

To investigate the possible involvement of DEK1 in regulation of cellulose biosynthesis or CWI signaling, WT and *DEK1* modulated lines were treated with Isoxaben. Isoxaben is a plant-specific herbicide which inhibits the synthesis of cellulose through interacting with CESAs and causing them to be removed from the PM and internalized into the cell (Heim et al., 1990; Shim et al., 2018). Isoxaben-induced disruption of cellulose synthesis causes reduced root growth and cell swelling and cellulose deficient mutants have been shown to display altered responses to Isoxaben treatment (Engelsdorf and Hamann, 2014; Vaahtera et al., 2019). We measured root lengths of 6-day old WT and *DEK1* modulated plants grown on media supplemented with either 2 nM Isoxaben or DMSO as a control (Figure 1A). In response to Isoxaben treatment WT plants showed a significant reduction of root length (Figure 1B). The mean root length difference between WT Isoxaben /DMSO (control) plants was −1.27 mm (95% CI −2.39, −0.0585, *p* = 0.0332). For DMSO control plants no differences in root lengths between WT and *OE CALPAIN* seedlings were observed. In contrast, we observed significant differences between the mean root length of WT and *OE CALPAIN* when grown on Isoxaben, shown by the 95% CIs of the mean differences of these two groups not overlapping (Figure 1B). Estimation statistics analysis showed high mean differences of −4.63 mm (95% CI −5.35, −3.95, *p* = <0.0001) in mean root length between *OE CALPAIN* plants grown on 2 nM Isoxaben supplemented media and the DMSO supplemented media controls (Figure 1B).

**Figure 1.**
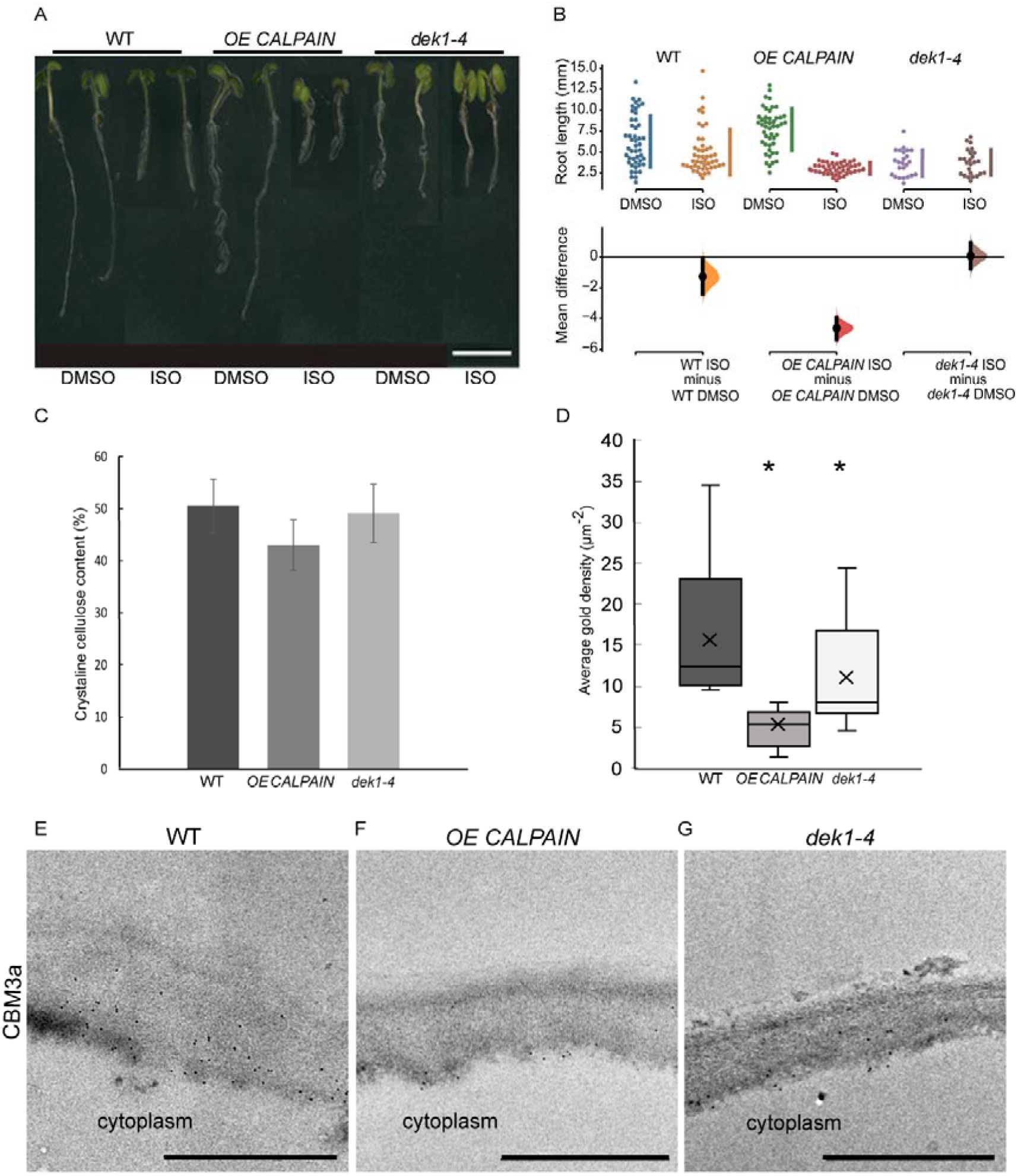
Isoxaben treatment and cell wall analysis of WT and *DEK1* modulated *Arabidopsis* lines. (**A**) Representative images of 6-day old WT, *OE CALPAIN* and *dek1-4* seedlings grown on media with either DMSO (control) or 2 nM Isoxaben, scale bar = 5 mm. (**B**) Multiple two-group estimation analysis of root length for each genotype between plants grown on media with DMSO and Isoxaben. The mean differences are shown in the Cumming estimation plot (lower panel). The raw data are plotted on the upper axes; each mean difference is plotted on the lower axes as a bootstrap sampling distribution. Mean differences are depicted as dots; 95% confidence intervals are indicated by the ends of the vertical error bars. N=47 WT DMSO, WT ISO, *OE CALPAIN* DMSO, 50 *OE CALPAIN* ISO, 24 dek1-4 DMSO, 25 dek1-4 ISO (**C**) Crystalline cellulose contents of AIR cell wall preparations of 10-day old seedlings. Analysis performed on two biological replicates and two technical replicates. Error bars are representing standard error. No statistical significance observed *p>0.05*, Student t-test. (**D**) Average gold particle density for CBM3a labelled cellulose. Asterix represents statistical significance, *p*<0.0001 for *OE CALPAIN, p*<0.05 for *dek1-4*, unpaired Student t-test. n=36 WT, 25 *OE CALPAIN* and 16 *dek1-4* epidermal cell walls. Analysis performed on biological duplicates. (**E-G**) Transmission electron micrographs of outer epidermal cell walls of 10-day old cotyledons showing immunogold labelled CBM3a cellulose binding. Scale bars = 500 nm.

In contrast, *dek1-4* mutants were completely insensitive to Isoxaben-induced mechanical perturbation of the cell wall (Figure 1A and 1B). Mean difference in the root length between plants grown on DMSO control media and plants grown on 2 nM Isoxaben was 0.0824 mm (95% CI −0.762, 0.94, *p* = 0.848). Since the mean difference in root length between untreated and Isoxaben treated *dek1-4* plants was almost 0, this indicated that Isoxaben treatment had no effect on *dek1-4* mutants. In addition, no significant differences in the mean difference were observed between *dek1-4* and WT (Figure 1B, 95% CI-s are overlapping).

### DEK1 is involved in fine tuning of epidermal cell wall composition

Studies of *DEK1* modulated lines grown on Isoxaben suggest cellulose content and/or assembly is altered. Cell wall linkage analysis was undertaken on alcohol insoluble residue (AIR) wall preparations to investigate potential wall compositional differences in 10-day old seedlings of *DEK1* modulated lines compared to WT. No statistically significant differences were found (Supplementary Figure 1, Supplementary Table 1). A trend towards reduced levels of cellulose in *OE CALPAIN* lines was observed and this was investigated further using the Updegraff (1969) method which determines crystalline cellulose content. Of the AIR, crystalline cellulose constituted 51% (w/w) in WT, 42% in *OE CALPAIN* and 48% in *dek1-4* (Figure 1C). Previous studies have shown DEK1-related changes in cellulose and pectin content predominantly occurs in the outer epidermal cell walls of both young and mature leaves and inflorescence stems (Amanda et al., 2016; Amanda et al., 2017). Immunofluorescence and immuno-gold labelling of cell wall epitopes was used to determine if wall composition differs at the tissue/cellular scale that may be beyond the resolution of the whole seedling chemical analyses. CBM3a labelling of cellulose epitopes showed slightly lower fluorescence intensity in the outer periclinal epidermal cell wall of both *DEK1* modulated lines compared to WT (Supplementary Figure 2A-C). TEM imaging of the cotyledon epidermal cell walls showed significantly decreased immuno-gold labelling of cellulose epitopes with CBM3a in both *DEK1* modulated lines compared with WT (Figure 1D-G). Average immuno-gold particle density for CBM3a in outer periclinal epidermal walls of 10-day old cotyledons was 14.09±1.01 μm^-2^ gold particles in WT, 5.93±1.01 μm^-2^ (*p*<0.0001) in *OE CALPAIN* and 10.41±1.44 μm^-2^ (*p*<0.05) in *dek1-4* (Figure 1D). Cotyledon sections labelled with Calcofluor white to detect cellulose, JIM5 and JIM7 to detect partially and highly methyl-esterified HG pectin, respectively, and LM15 to detect XG revealed no obvious differences in fluorescence signal between WT and *DEK1* modulated lines (Supplementary Figure 3). We also performed immuno-gold labelling with JIM5 and JIM7 antibodies that detect HG pectins in cotyledon epidermal cell walls (Supplementary Figure 4). Our results show that *dek1-4* had significantly higher average density of JIM5 immuno-gold particles (26.73±1.91 μm^-2^, *p*<0.0001) compared with WT (18.86±0.82 μm^-2^), while *OE CALPAIN* has a significantly lower count of JIM5 immuno-gold particles (15.79±1.10 μm^-2^, *p*=0.02) (Supplementary Figure 4A-C and 3G). Levels of partially-methyl esterified HG detected by the JIM7 antibody were significantly higher in *OE CALPAIN* (3.35±0.24 particles μm^-2^, *p*=0.03) and significantly lower in *dek1-4* (1.90±0.21 μm^-2^, *p*=0.01) compared to WT (2.63±0.19 μm^-2^) (Supplementary Figure 4D-F and 3H).

### CESA transcript levels show no differences between WT and *DEK1* modulated lines

Isoxaben growth assays, immuno-labeling and cell wall compositional analysis suggest DEK1 influences levels of cellulose, particularly in epidermal walls. To determine how changes in cellulose may have come about we investigated CSCs at both the transcriptional and post-transcriptional levels. Previous studies have shown that modulation of DEK1 affects expression of numerous cell wall-related genes, however *CESA* levels were unaffected (Amanda et al., 2016; Johnson et al., 2008). *CESA* transcript levels were investigated in 6-day old seedlings of WT, *OE CALPAIN* and *dek1-4* lines using qRT-PCR. No significant differences in transcript levels of *CESA1, CESA3* and *CESA6*, encoding primary wall CESAs, were observed between WT and *DEK1* modulated lines (*p*>0.05) using fold change analysis (Supplementary Figure 5). These data suggest DEK1 influences cellulose biosynthesis at the post-transcriptional level.

### *DEK1* modulated lines influence mobility and not density of CSCs

Changes in cellulose content could be explained by differences in the number of CSCs at the PM. *OE CALPAIN* and *dek1-4* lines were crossed with a pCESA3:GFP:CESA3 mCh:TUA5 (GFP:CESA3) reporter line and the number of GFP:CESA3 complexes on pavement cell PM patches quantified (Figure 2A). The average density of CSC particles in WT was 0.89 ± 0.04 particles μm^-2^, *OE CALPAIN* was 0.93 ± 0.04 particles μm^-2^ and *dek1-4* had 0.82 ± 0.04 particles μm^-2^. Thus, no statistically significant differences were detected in CSC particle density between WT and *DEK1* modulated lines (Figure 2B, Students t-test, *p*>0.05). We then proceeded to investigate if DEK1 could affect the velocity of the CSC. Investigation of CSC velocity is commonly used as a proxy for determining cellulose synthesis rates at the cellular level. In brief, the more cellulose CESAs synthesize, the faster CSCs will move along within the plane of the PM (Paredez et al., 2006; Diotallevi and Mulder, 2007; Fujita et al., 2011). CSCs at the PM of the periclinal epidermal cells of 3-day old cotyledons were tracked during 10-min time lapses (Figure 2C) and kymographs of GFP-labelled CESA3 particles (Figure 2D) were used to obtain velocities of CSCs (Figure 2E). CSC velocities in both *OE CALPAIN* and *dek1-4* mutant lines were significantly reduced compared to WT (Figure 2E). WT CSCs had an average velocity of 192.41 ± 4.16 nm min^-1^, *OE CALPAIN* 124.11 ± 3.01 nm min^-1^ and *dek1-4* 113.91 ± 3.09 nm min^-1^. The reduction in CSC velocity in both *DEK1* modulated lines was highly statistically significant compared to WT (Unpaired Student t-test, *p*<0.0001). A reduction of CSC velocities in the periclinal cell walls of cotyledon pavement cells is consistent with the findings of immuno-histochemical assays that indicate significantly reduced cellulose levels in *OE CALPAIN* compared to WT (Figure 1D).

**Figure 2.**
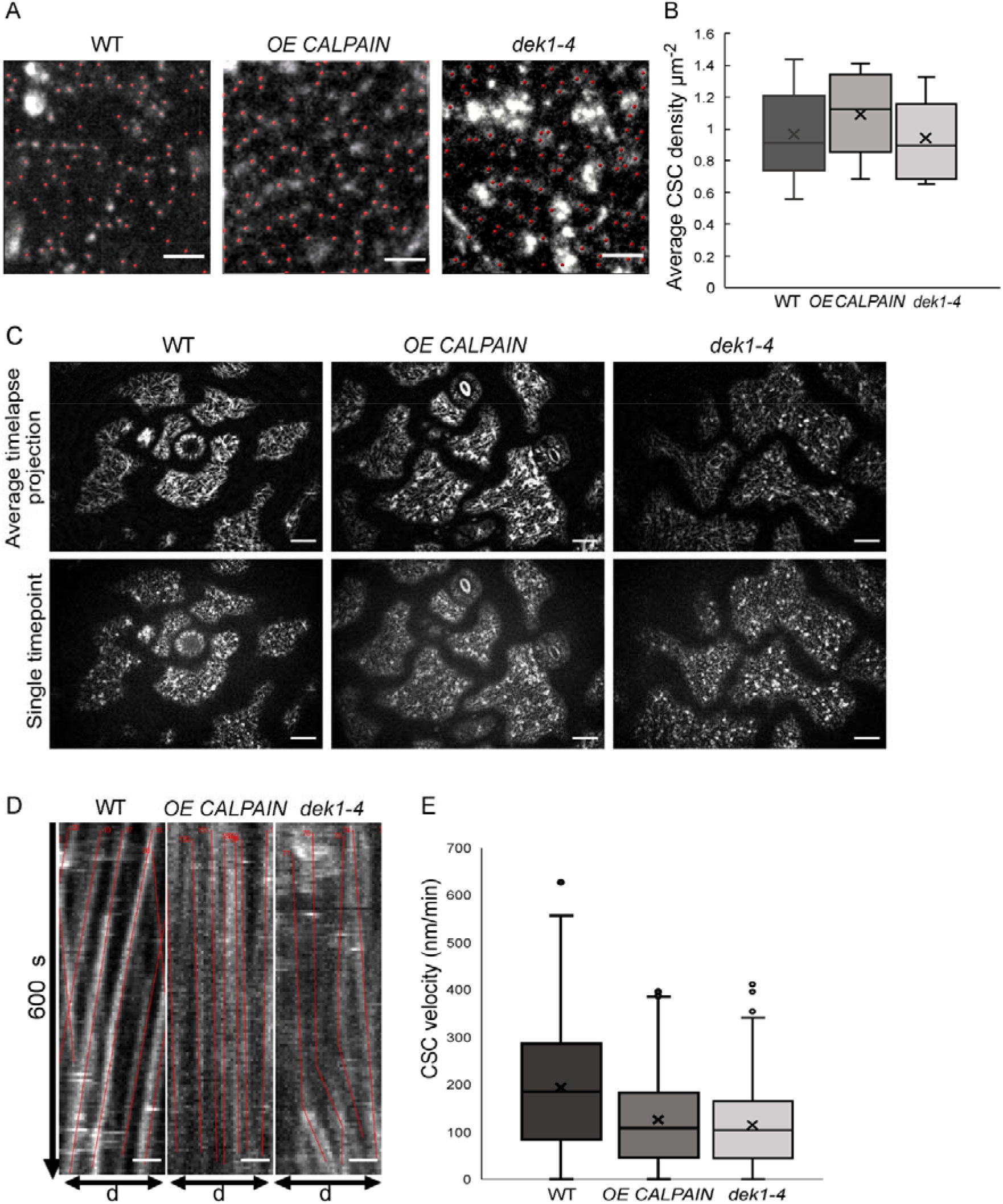
Analysis of cellulose synthase complex (CSC) migration at the plasma membrane in WT and DEK1 modulated *Arabidopsis* lines. (**A**) Representative micrographs of 3-day old *Arabidopsis* cotyledon epidermal pavement cells’ membrane patches expressing GFP-CESA3 fluorescent marker used for CSC (red dots) density calculation in Col-0 and *DEK1* modulated lines. Scale bars = 2 μm. (**B**) Box plots showing average density of CESA particles in 3-day old cotyledon pavement cells. n = 6 plants for WT, 31 cells; 6 plants for *OE CALPAIN*, 33 cells; 6 plants for *dek1-4*, 27 cells. No statistical significance observed between WT and *DEK1* modulated lines, *p*=0.5647 for *OE CALPAIN, p*=0.2705 for *dek1-4*. (**C**) Representative images of 3-day old cotyledon epidermal pavement cells used for generation of kymographs. Upper panel represents average projections of time lapses, bottom panel represents a single frame of a corresponding time lapse, n=6 plants per genotype. Scale bars = 10 μm. (**D**) Representative kymographs showing migration of CESA complexes in 3-day old cotyledon pavement cells of Col-0 and *DEK1* modulated lines. Red lines represent trajectories of individual CESA particles. WT, n=6 plants, 937 CSC particles tracked; *OE CALPAIN*, n=6 plants, 982 CSC particles tracked; *dek1-4*, n=6 plants, 692 CSC particles tracked. Analysis performed on biological duplicates. Scale bars = 10 μm. (**E**) Box plots represent distribution of CESA velocities, obtained from kymographs. Asterisk represent statistical significance (*p*<0.0001, Student t-test).

### CSC membrane trafficking is significantly altered in *DEK1* modulated lines

To investigate if DEK1 might be regulating other functional properties of CSC complexes, delivery rate of CSCs and their lifetime at the PM was examined. Fluorescence recovery after photobleaching (FRAP) experiments were performed on *OE CALPAIN* and *dek1-4* plants crossed into a GFP:CESA3 reporter line to determine if the rate of exocytosis of CSCs into the PM was altered. Imaging of GFP:CESA3 particles during the 10 min following photobleaching (Figure 3A) showed CESAs in *OE CALPAIN* had the fastest recovery after FRAP (Figures 3B and 3C) with 3.96 ± 0.36 CSC μm^-2^h^-1^ (*p*= 0.0188) compared to WT with a CSC recovery rate of 2.86 ± 0.25 CSC μm^-2^h^-1^. The lowest CSC recovery rate post-FRAP was observed in *dek1-4* (Figures 3B and 3C) with only 1.87 ± 0.15 CSC μm^-2^ (*p*=0.0033). Studies of the lifetime of CSC particles at the PM showed the opposite trends (Figure 3D). *OE CALPAIN* CSCs spent an average of 14.33 ± 4.69 min in the PM, which was significantly shorter than WT CSCs with 19.56 ± 1.42 min (*p*=0.0288), while *dek1-4* CSCs stayed significantly longer in the PM (25.36 ± 1.73 min, *p*=0.0152) before undergoing endocytosis (Figure 3D).

**Figure 3.**
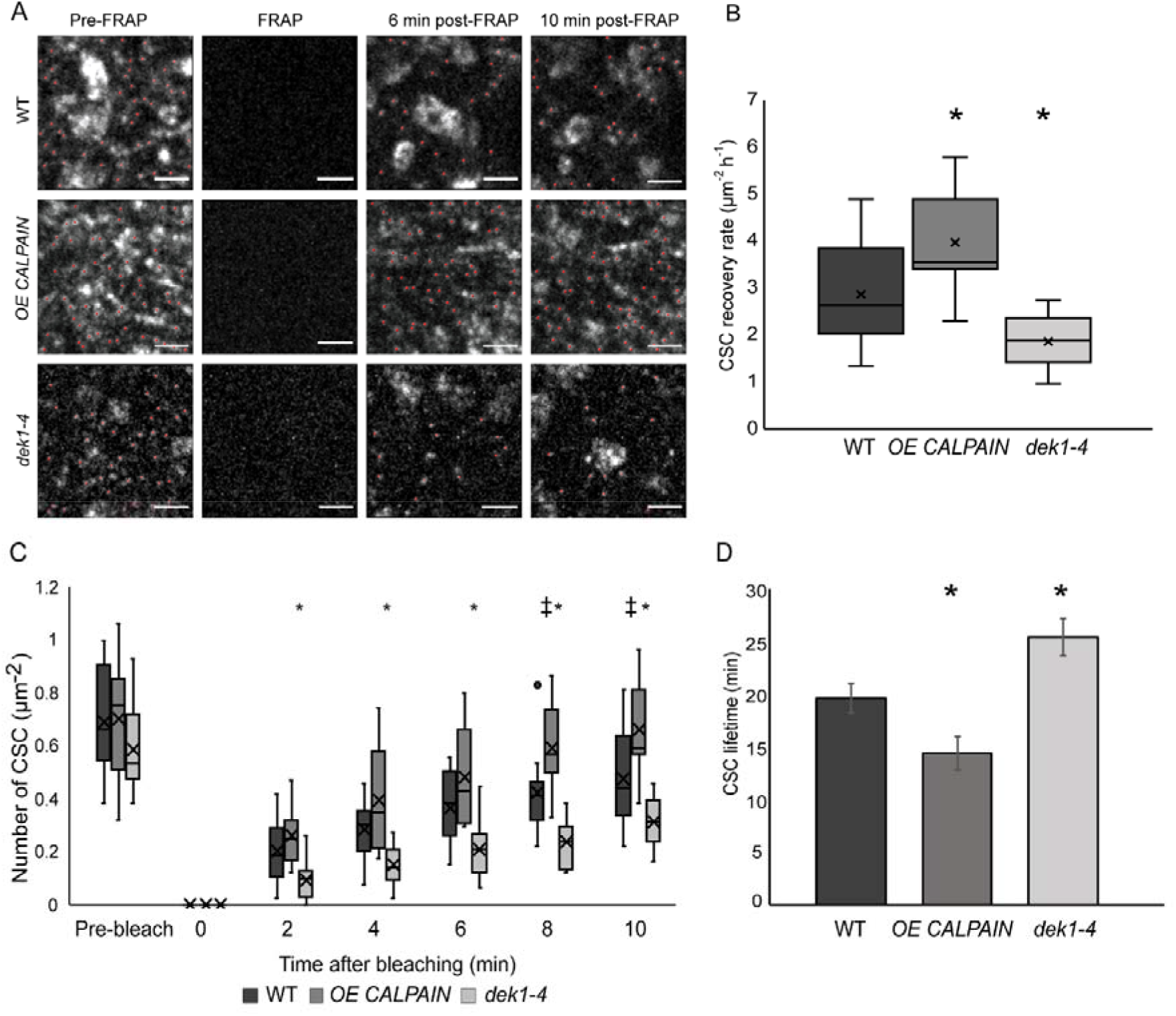
Delivery and resident time of CESAs at the plasma membrane of WT and *DEK1* modulated lines in *Arabidopsis* using fluorescent-recovery after photo-bleaching (FRAP). (**A**) Representative images showing delivery of new CESA complexes (red dots) after photo-bleaching in 3-day old cotyledon pavement cells of WT, *OE CALPAIN, dek1-4*. Scale bars = 2 μm. (**B**) Box plots showing average recovery rate of CESA particles following FRAP in cotyledon pavement cells. Asterisk represent statistical significance (*p*<0.0188 for *OE CALPAIN, p*<0.0033 for *dek1-4*, Student’s t-test). (**C**) Box plots representing recovery of CESA particles after photo-bleaching at each time-point. Symbols above box plots represent statistical significance (*p*=0.0102 and 0.0188 for *OE CALPAIN, p=*0.0001-0.0055 for *dek1-4*, Student’s t-test). (**D**) Histogram representing average lifetime of CESA complexes at the PM. Asterisk above boxplot represents statistical significance (*p*<0.05, Student’s t-test), error bars represent standard error. (**B-D**) Analysis was performed in two biological replicates for WT and *dek1-4*, and one replicate for *OE CALPAIN*. n = 7 cotyledons for WT, 3 cotyledons for *OE CALPAIN*, 6 cotyledons for *dek1-4;* n = 16 cells for WT, 9 cells for *OE CALPAIN*, 14 cells for *dek1-4*.

### DEK1 affects cell wall mechanics of cotyledon pavement cells

The defects in CESA dynamics observed in cotyledon pavement cells prompted us to examine the possible role of DEK1 in influencing structure and physical properties of the CMFs and consequently the mechanical properties of the cell wall. We used atomic force microscopy (AFM) to image and mechanically characterize native cell walls.

We performed AFM imaging on dry cell wall monolayers of isolated abaxial epidermal cell walls from cotyledons of 10-day old *Arabidopsis* seedlings (Figure 4A-C; Supplementary Figures 6 and 7). Imaging was done on the inner surface of the abaxial epidermis outer periclinal walls, which is the newest deposited layer of the cell wall, abutting the PM (Supplementary Figure 6). Cell wall images of WT clearly show the CMF network (Figure 4A-C, Supplementary Figure 8), as reported for onion epidermal cell wall monolayers imaged by AFM (Kafle et al., 2014; Zhang et al., 2016). Differences in the thickness of CMFs were observed in *OE CALPAIN* and *dek1-4* (Figure 4A-C) compared to WT. In *OE CALPAIN*, thicker bundles of CMFs were observed as well as amorphous deposits on the cell wall surface (Figure 4B). Diameters of the CMF bundles were quantified from height channel images (Supplementary Figure 8). Results showed that the CMF bundles of both *OE CALPAIN* and *dek1-4* had higher mean diameters than WT CMFs (Figure 4D). Mean diameter of WT CMFs was 11.09 ± 0.03 nm. For *OE CALPAIN* CMF it was 16.21 ± 0.03 nm, and for *dek1-4* it was 15.74 ± 0.03 nm. These differences are highly statistically significant (*p* <0.0001). Analysis of the distribution of CMF dimeters shows that both *DEK1* modulated lines have a higher frequency of bundled CMFs in the range of 15-40 nm whereas WT CMFs are mostly represented in the range of 1-15 nm, implying that WT has either an abundance of single CMFs (~3.5 nm is the diameter of a typical CMF (Zhang et al., 2016)) or smaller bundles of CMFs (Figure 4E).

**Figure 4.**
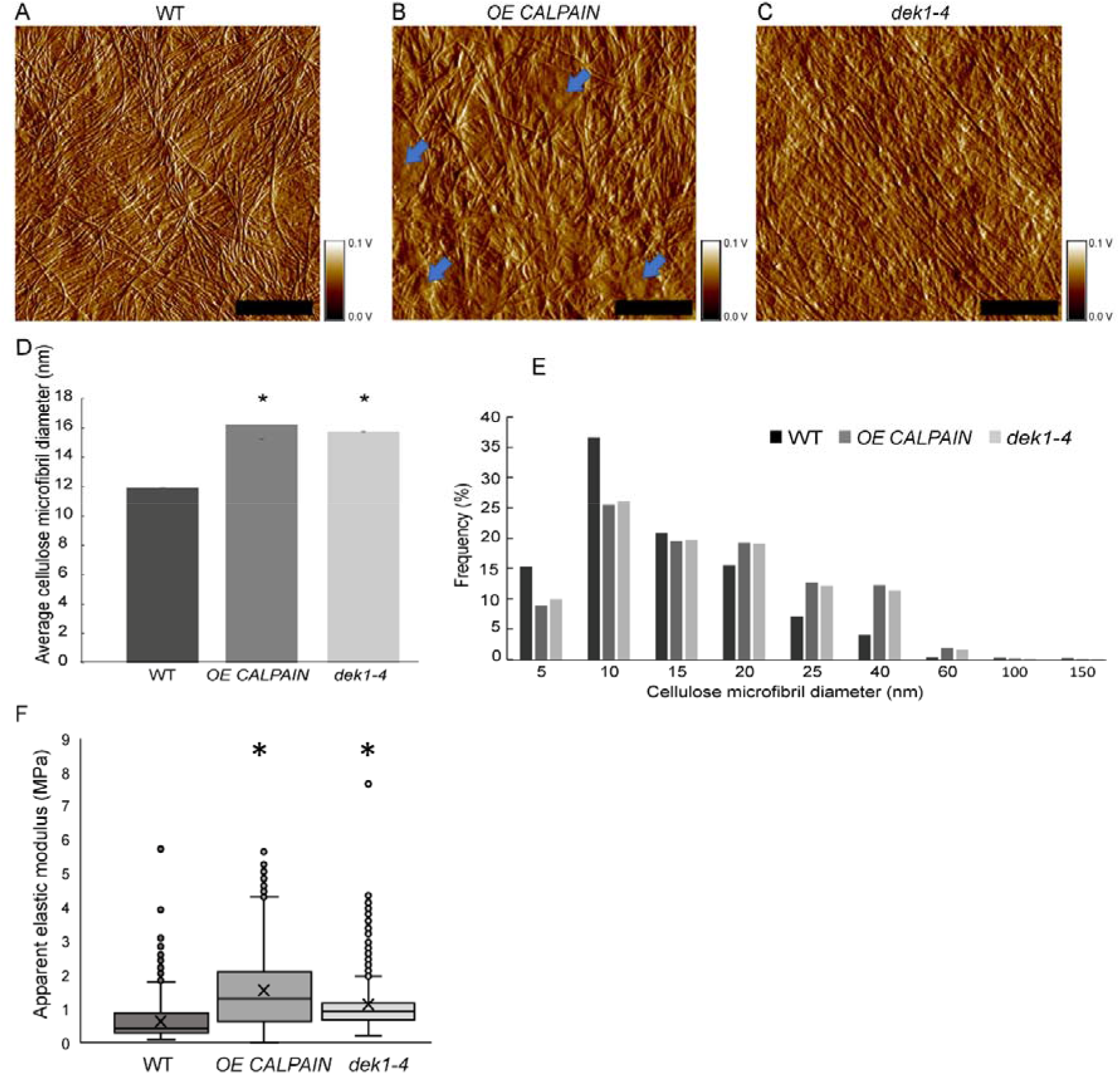
AFM imaging and and nanomechanical characterization of outer periclinal cell walls. (**A-C**) AFM amplitude channel images of the inner face (juxtaposed to the PM) of periclinal primary cell walls in abaxial cotyledon pavement cells of 10-day old seedlings of WT, *OE CALPAIN* and *dek1-4*. Samples were imaged using tapping mode in air. Arrows indicate amorphous cell wall regions. Scale bars = 250 nm. Corresponding height channel AFM images are shown in Supplementary Figure 8. (**D**) Histograms representing average thickness of cellulose microfibrils in cotyledon epidermal cell walls. Asterisk represents statistical significance, *p*<0.0001, unpaired Student t-test. n=5 cotyledons, 10 cells, 1-2 regions per cell for WT; n=6 plants, 6 cells, 1-2 regions per cell for *OE CALPAIN;* n=4 cotyledons, 7 cells, 1-3 regions per cell for *dek1-4*. (**E**) Distribution of CMF thickness and frequency in WT, *OE CALPAIN* and *dek1-4*. (**F**) Box plots representing distribution of cell wall apparent elastic moduli generated after nanoindentation of extracted cotyledon epidermal cell walls in a direction perpendicular to CMFs of 10-day old cotyledons of WT, *OE CALPAIN* and *dek1-4*. n=12 indented cotyledons from 12 different plants for WT (1427 indentation points), 9 cotyledons from 9 different plants for *OE CALPAIN* (1553 indentation points), and 12 cotyledons from 12 different plants for *dek1-4* (1509 indentation points). One cell per one cotyledon was indented in all genotypes. Asterisk represents statistical significance, *p*<0.0001, Student t-test. (**A-F**) All experiments were performed in two biological replicates.

Previous studies have shown that changes of cell wall composition in cell wall mutants of *Arabidopsis* can affect mechanical behaviour of the cell wall and CMF properties (Xiao and Anderson, 2016). To examine cell wall mechanical properties in *DEK1* modulated lines, we performed AFM nanoindentation on the isolated abaxial epidermal cell walls of 10-day old *Arabidopsis* cotyledons. Nanoindentation generated force-displacement curves, which were used to calculate cell wall elastic moduli (Supplementary Figure 9A, C, E). It is worth noting that within all analyzed genotypes, a large range of elastic moduli was observed (Figure 4F, Supplementary Figure 9B, D, F). In WT, values ranged from 0.1 MPa to 5 MPa, which is consistent with previous data showing highly heterogenous distribution of cell wall mechanical properties (Yakubov et al., 2016). Mean apparent elastic modulus of WT was 0.71 ± 0.01 MPa. Analysis of the apparent elastic modulus in *OE CALPAIN* and *dek1-4* suggests that both lines had significantly stiffer cell walls compared to WT. *OE CALPAIN* had an elastic modulus of 1.52 ± 0.03 MPa, while *dek1-4* had an elastic modulus of 1.11 ± 0.02 MPa (Figure 4F). These results suggest DEK1 is involved in regulating and maintaining mechanical properties of the cell wall.

### Stiffer cell walls of *OE CALPAIN* and *dek1-4* exhibit absence of ectopic lignification

The altered cell wall properties in DEK1 modulated lines, including cellulose levels, CMF thickness and cell wall mechanics along with changes in CESA dynamics led us to investigate if these changes could be associated with disrupted CWI sensing/signalling pathways. One of the regularly observed responses from either cell wall damage or disrupted cell wall synthesis, caused by either abiotic or biotic factors is ectopic deposition of lignin. WT and DEK1 modulated 6-day old seedlings were grown overnight in liquid MS media supplemented with either DMSO control or 600 nM Isoxaben and then stained with phloroglucinol to detect lignin. WT developed strong lignin staining in the elongation region behind the root tip in response to Isoxaben treatment (Figure 5A and B), while both *DEK1* modulated lines developed weak, barely detectable lignin staining (Figure 5A and B).

**Figure 5.**
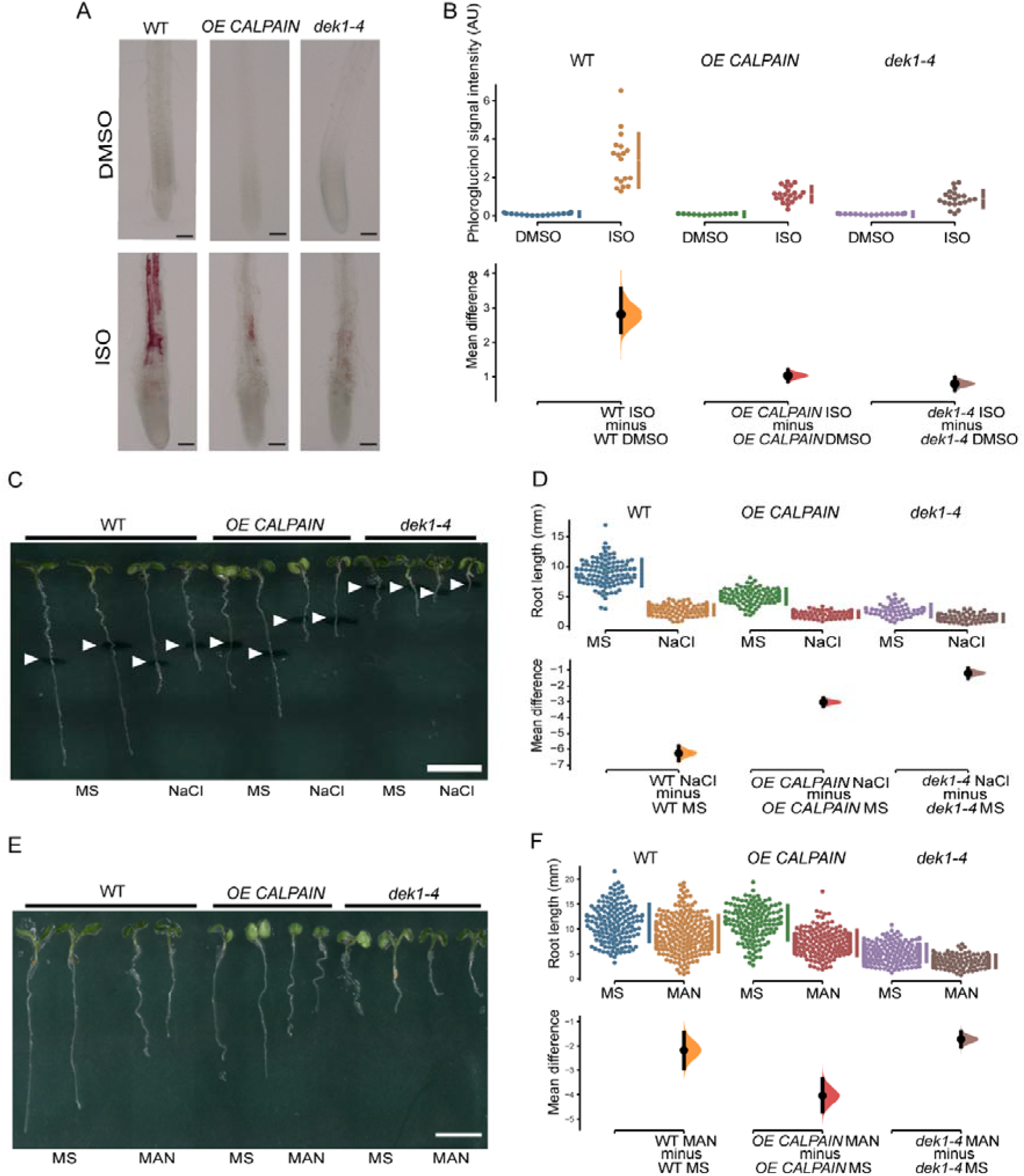
Response of WT and *DEK1* modulated *Arabidopsis* lines to cellulose perturbation, salt and osmotic stress. (**A**) Phloroglucinol staining of Isoxaben-induced ectopic lignin deposition in elongation region near root tips of 6-day old WT, *OE CALPAIN* and *dek1-4* seedlings grown in media with either DMSO (control) or 600 nM Isoxaben (ISO). Scale bars = 100 μm. (**B**) Mean difference for arbitrary intensity values of phloroglucinol signal from roots in (**A**) are shown in the Cumming estimation plot. N = 16 WT DMSO, 18 WT ISO, 12 *OE CALPAIN* DMSO, 21 *OE CALPAIN* ISO, 18 *dek1-4* DMSO, 21 *dek1-4* ISO (**C**) Representative images of 7-day old WT, *OE CALPAIN* and *dek1-4* seedlings transferred for 48 h on media supplemented with or without 140 mM NaCl. White arrowheads indicate length of roots at transfer. Scale bar = 5 mm. (**D**). Mean difference for root length for each genotype between plants grown on control media and plants grown on NaCl supplemented media. N = 107 WT MS, 108 WT NaCl, 95 *OE CALPAIN* MS, 97 *OE CALPAIN* NaCl, 53 *dek1-4* MS, 76 *dek1-4* NaCl. (E) Representative image of light-grown 6-day old WT, *OE CALPAIN* and *dek1-4* seedlings grown on control media (MS) or media supplemented with 200 mM mannitol (MAN). Scale bar = 5 mm. (**F**) Mean difference for root length for each genotype between plants grown on MS media or media with mannitol. N = 152 WT MS, 230 WT MAN, 140 *OE CALPAIN* MS, 181 *OE CALPAIN* MAN, 235 *dek1-4* MS, 133 *dek1-4* MAN. (**B,D,F**) Mean difference for lignin staining intensity and root length under different experimental conditions are shown in the Cumming estimation plot. The raw data are plotted on the upper axes; each mean difference is plotted on the lower axes as a bootstrap sampling distribution. Mean differences are depicted as dots on the lower panels of Cumming plots; 95% confidence intervals are indicated by the ends of the vertical error bars.

### *OE CALPAIN* and *dek1-4* show altered responses to salt and osmotic stress

To determine the response of *DEK1* modulated lines to other types of cell wall damage, salt treatment was used to disrupt pectin HG Ca^2+^ cross-linking in cell walls. Transfer of 5-day old plants from control MS media to MS media supplemented with 140 mM NaCl for 48 h resulted in severe inhibition of root growth of WT (Figure 5C). Mean difference of root growth between WT grown on control media and plants transferred to NaCl supplemented media was −6.25 mm (95% CI −6.71, −5.81, *p*<0.0001 two-sided permutation t-test). The response of *OE CALPAIN* plants to NaCl was more moderate (Figure 5C and D) with a root growth mean difference between control and salt-grown plants of −3.02 mm (95% CI −3.29, −2.75, *p*<0.0001 two-sided permutation t-test). A similar insensitive response to that of Isoxaben treatment was observed for *dek1-4* NaCl treatment (compare Figures 1B and 5D), with a small mean difference between control and NaCl-grown plants (Figure 5D). In general, both *DEK1* modulated lines showed reduced sensitivity to NaCl when compared to WT (Figure 5C and 5D).

CWI is also affected by changes in osmotic pressure of the cell, which causes either protoplast shrinking or swelling depending on whether the osmotic pressure drops or rises, respectively, in the protoplast. Both types of changes have an impact on cell wall composition and its integrity (Vaahtera et al., 2019). This led us to investigate osmotic responses in WT and *DEK1* modulated lines by growing them on 200 mM mannitol supplemented MS media. Response to mannitol treatment in 6-day old light grown plants was shown to have a similar trend to Isoxaben-induced cell wall perturbations (Figure 5E). Mannitol treatment had very little effect on WT (Figure 5F), with a root length mean difference between mannitol- and MS-grown plants of −2.18 mm (95% CI −2.95, −1.45, *p*<0.0001). In contrast, *OE CALPAIN* plants showed higher sensitivity to mannitol treatment with root length mean difference between mannitol treated and control grown plants of −4.05 mm (95% CI −4.7, −3.36, *p*<0.0001), that is, almost double the decrease in root length of treated/control plants compared with WT (Figure 5F). Finally, *dek1-4* roots showed decreased sensitivity to mannitol, similar to that observed in Isoxaben and salt assays. Root length mean difference between mannitol and control treated *dek1-4* plants was −1.73 mm (95% CI −2.03, −1.42, *p*<0.0001). The effect of mannitol on root growth of *OE CALPAIN* plants was statistically significant in both WT and *dek1-4*, since there was no overlapping of *OE CALPAIN* root length 95% CI with root length CIs of both WT and *dek1-4* (Figure 5F). In contrast, both WT and *dek1-4* had no statistical differences in response to mannitol treatment since their root length 95% CI overlap (Figure 5F).

In conclusion, DEK1 affects the synthesis of both cellulose and pectins in early developmental stages of *Arabidopsis*. Both *DEK1* modulated lines show decreased crystalline cellulose in outer epidermal cell wall of cotyledons. Velocity of CSCs in the epidermal pavement cell PMs, which is used as a proxy for the biosynthetic activity, is significantly reduced in both *DEK1* modulated lines, while the density of CSC complexes is similar to that of the WT. PM dynamics of CSCs is significantly altered in both lines, with *OE CALPAIN* CSCs having significantly slower CSC exocytosis and faster endocytosis, while these features of CSC dynamics were reversed in *dek1-4* plants. Epidermal cell wall stiffness was significantly increased in both *DEK1* modulated lines compared to WT.In addition, *DEK1* modulated lines exhibited changes in their ability to detect changes in CWI, most notably hypersensitivty of *OE CALPAIN* and complete insensitivity of *dek1-4* to Isoxaben-induced inhibition of cellulose synthesis and ectopic lignification.

## DISCUSSION

### DEK1 in cellulose assembly

DEK1 is proposed to act as a mechano-sensor protein at the PM (Amanda et al., 2016; Tran et al., 2017). Upon perception of mechanical stimuli, originating either from other growing cells in the tissue or from a cell’s own turgor pressure, or an external environmental signal, the CALPAIN domain of DEK1 is proposed to be auto-catalytically released into the cytoplasm (Amanda et al., 2016; Tran et al., 2017). DEK1 CALPAIN likely acts as a regulatory protease (Wang et al., 2003; Johnson et al., 2008) and downstream signalling initiates changes in cell wall biosynthesis/remodelling and epidermal specification (Demko et al., 2014; Amanda et al., 2016; Amanda et al., 2017; Galletti et al., 2015). In this study we show DEK1 influences CESA dynamics, cellulose content, pectin re-structuring and mechanical properties of epidermal walls, and propose a role in CWI sensing.

It is currently understood that slower movement of CSCs across the PM, a proxy for a lower rate of cellulose biosynthesis, results in less cellulose synthesized (Diotallevi and Mulder, 2007). The significant reduction of CSC velocity in *OE CALPAIN* and *dek1-4* (Figure 2D and E) support a role for DEK1 in regulation of CESA dynamics. Mutants for *CESA6/prc1-1*, and *CESA1/any1*, have significantly reduced CESA velocities in dark grown hypocotyls (Bischoff et al., 2011) that is proposed to be the major cause of a 30% reduction in growth (Fagard et al., 2000). There is no data in the literature that CESA proteins undergo proteolytic processing after insertion (by exocytosis) into the PM. RT-qPCR analysis of *CESA1,3* and *6* expression levels, and measurement of CESA density at the cotyledon pavement epidermal cell PMs showed no difference between WT and *DEK1* modulated lines. These data support DEK1 regulation of CSCs at the post-translational level. An interesting possibility is that DEK1 regulates CSC activity indirectly, potentially through interactions with either CESA regulatory proteins or the cytoskeleton components that guide CSCs. To more precisely define the mechanism, crosses of *DEK1* modulated lines to mutants of different CESA regulatory proteins as well as with their reporter lines should be undertaken. Candidates would include CSI1/POM2 that facilitate binding between CSCs and CMTs (Gu et al., 2010; Bringmann et al., 2012), the endo-1,4-beta-glucanase KORRIGAN (KOR) involved in CESA regulation (His et al., 2001; Vain et al., 2014) and PATROL1 (PTL1) that interacts with exocyst complex proteins and CS1/POM2 to deliver CSCs to the PM (Zhu et al., 2018). Mutants *kor1-1* and *kor1-3* display a similar reduction in CSC mobility to *DEK1* modulated lines when compared to WT plants (Vain et al., 2014). In addition *kor* mutants have been shown to have decreased cellulose content, but also increased pectin abundance, which was interpreted as a compensatory mechanism and part of feedback loops connecting cellulose and pectin synthesis to changes in the mechanical properties of the cell wall (His et al., 2001).

It could be conceivable that DEK1 is involved in CSC guidance along CMTs, either by modifying the activity and properties of CSI1/POM2 or some other CESA-CMT interface proteins. Reduced CSCs velocities and cellulose content is observed in *pom2-1* mutants compared to WT plants as well as spiral twisting of the entire rosette and leaves. Previous studies of lines with reduced levels/activity of *DEK1*, including *dek1-4*, showed epinastic curling of cotyledons and leaves (Roeder et al., 2012; Galletti et al., 2015). CMTs are responsive to mechanical stress and assumed to align in the direction of maximal stress (Hamant et al., 2008). CMTs were shown to re-organize normally in *dek1-4* mutants in response to ablation in cotyledons, however, were more highly orientated than in WT pavement cells due to changes in cell shape (Galletti et al., 2015). A study in animals has shown that a non-proteolytic variant of CALPAIN, CALPAIN6 (CAPN6), stabilizes microtubules by binding to them and preventing their de-polymerization (Tonami et al., 2011). Unlike CAPN6, DEK1 CALPAIN has proteolytic activity, and it would be interesting to investigate if DEK1 CALPAIN can interact with CMT and potentially disrupt binding of CESA complexes (Tonami et al., 2011).

Upregulation of the exocytotic module of the CSC trafficking machinery in *OE CALPAIN* (Figure 3B and C) could be one explanation for the faster exocytosis of the CESAs, as observed by faster post-FRAP recovery of CSC particles (Figure 3A and B). Exocytosis of CSCs is regulated by a large protein complex (Polko and Kieber, 2019), which interacts with CS1 and PTL1 to deliver CSCs to the PM (Zhu et al., 2018). The frequency of CSC insertions and alignment with the CMT was highly decreased and the delivery rate of CSC severely diminished in *csi1-3/ptl1-2* double mutants. In addition, *ptl* mutants also have lower CSC velocities, and both *ptl1-2* and *csi1-3/ptl1-2* lines have significant decreases in cellulose content (Zhu et al., 2018). The contrasting effect which *OE CALPAIN* and *dek1-4* have on CSC PM trafficking could mean that DEK1 is interacting with several components involved in CSC trafficking to (and in) the PM, and that its role in signalling involves multiple target proteins.

### DEK1 in CWI sensing

Isoxaben growth assays point to a possible role of DEK1 in CWI signalling and maintenance (Figure 1A and 1B). CWI maintenance systems are thought to be activated in response to reduced cellulose levels, such as induced by Isoxaben treatment (Denness et al., 2011; Hamann, 2014; Basu et al., 2016) to limit damage and growth. The hypersensitivity to Isoxaben in *OE CALPAIN* plants (Figure 1A and 1B) could be a combination of reduced cellulose and increased activation of CWI signaling as a result of excess CALPAIN levels. A slight trend towards reduced cellulose levels was also observed in *dek1-4* in which CALPAIN activity is modulated by a single nucleotide polymorphism (Roeder et al., 2012). The insensitivity to Isoxaben observed in *dek1-4* suggests CWI signaling is impaired and supports a role for DEK1 in the sensing/signaling pathways. DEK1 is likely a multifaceted regulatory protease, participating in different regulatory networks during early and late developmental stages and different tissue types. In *Arabidopsis* DEK1 could additionally initiate compensatory pathways as a result of cell wall changes as has been shown to occur in mutants with defects in cell wall biosynthesis (His et al., 2001; Hu et al., 2019).

The epidermis is a key regulator of plant growth and this is influenced by the mechanical properties of the cell wall (Galletti et al., 2016; Moulia et al., 2021). In *DEK1* modulated lines, pectins could act as a compensatory mechanism to counteract the decreased production of cellulose in order to preserve the CWI and lead to increased stiffness of the cell wall (His et al., 2001). DEK1 has previously been shown to regulate expression of pectin-related genes, including *GAUTs* and pectin methyltransferases as well as altered deposition of pectic polysaccharides in leaf epidermal cell walls of *OE CALPAIN GFP* (Amanda et al., 2016; Amanda et al., 2017). Mutants in pectin methyltransferase, *quasimodo2* and *tumorous shoot development2*, have decreased levels of both HGs and cellulose, decreased CESA velocities, thinner and more fragmented CMFs (Mouille et al., 2007; Du et al., 2020), while mutants in putative galacturonic acid transferase, *quasimodo1*, experience severe defects in cell-to-cell adhesion due to low HG-content (Bouton et al., 2002; Verger et al., 2018). Increased partially methyl-esterified HG in *dek1-4* is consistent with a role for pectin-related increased bundling of CMFs in this line. Stiffer cell walls in *OE CALPAIN*, with reduced levels of partially methyl-esterified HG and increased levels of highly methyl-esterified HG requires further investigation. The relationship between the degree of HG methyl-esterification and cell wall stiffness is unresolved. It could depend upon local concentrations of Ca^2+^ in the apoplast (Wang et al., 2020), orientation of the cell wall, such as anticlinal *vs* periclinal cell walls (Haas et al., 2020), activity of polygalacturonases (Yang et al., 2018) and many as yet unknown factors. Changes in interactions between pectins and CMF could also influence mechanical properties of the cell wall in *DEK1* modulated lines (Figure 4E). NMR studies of ‘never dried’ onion epidermal cell walls have shown that HGs make the majority of contacts with CMFs, which could mean that they act as primary mechanical tethers (Wang et al., 2015). Pectins affect cell wall mechanical properties through poro-elastic effects, without impacting the structural properties of cellulose (Lopez-Sanchez et al., 2015).

Regulation of cell wall biomechanics in response to mechanical stress and the potential influence on cell and plant morphogenesis has highlighted the importance of CWI. CWI pathways are now seen as key to modulating activity of wall biosynthesis/remodelling and growth. Previous studies have revealed ectopic lignification as an important element of CWI maintenance responses (Denness et al., 2011; Hamann, 2014; Vaahtera et al., 2019). Lack of ectopic lignification was expected in *dek1-4* since this line showed insensitivity to Isoxaben suggesting a reduced capacity to initiate CWI responses. The loss of ectopic lignification in *OE CALPAIN* was intriguing. It is possible that constitutive overexpression of CALPAIN, proposed to activate mechanical stress responses, has been dampened over-time by other regulatory pathways, limiting lignin production, however this needs further investigation.

Sensitivity of *DEK1* modulated lines to abiotic stresses such as salt and osmotic pressure (simulating drought) add further support to the potential role in CWI signalling (Figure 5C-F). Different stress conditions are known to lead to similar responses of plants in terms of growth and morphogenesis (Tenhaken, 2014). However, the underlying molecular mechanism of these changes and their associated signalling cascades can differ greatly. CWI sensors of the CrRLK family, including FER and THE1 are involved in responses to stress, mediated by RALF peptide binding (Feng et al., 2018; Blackburn et al., 2020; Zhang et al., 2020). RALF peptides are known to be processed by substilin-like proteases (Stegmann et al., 2017), however, other proteases such as DEK1 may be involved in generating mature RALF ligands. DEK1 signalling cascades are still completely unknown and further work is required to determine if cross-talk between CrRLK signalling and DEK1 pathways occur. Identification of CALPAIN targets through the use of targetted proteomics approaches is essential for understanding the pathways through which DEK1 acts, and its precise role in plant growth and development, including CWI.

## MATERIAL AND METHODS

### Plant resources

*Arabidopsis thaliana* Columbia-0 (Col-0) ecotype was used as wild type (WT) control. *dek1-4* is an EMS single point mutant (Roeder et al., 2012) backcrossed into Col-0 as described in (Galletti et al., 2015). The CALPAIN overexpression construct *pRPS5A:CALPAIN:6X HISTIDINE* (*OE CALPAIN*), with hygromycin resistance in Col-0 wild type background is described in Galletti et al., 2015. For CSC dynamics analysis we used *OE CALPAIN* and *dek1-4* crossed into CSC reporter line *pCESA3:GFP:CESA3* (Desprez et al., 2007).

Plants were grown vertically in petri dishes, on MS 1% (w/w) sucrose media, 0.8% agar, pH 5.8 (Murashige and Skoog, 1962), without vitamins. Plants were grown in Percival growth cabinets, model CU36L/LT (Percival Scientific, Perry, Iowa, USA), at a temperature of 21°C and light intensity of 95-111 mol m^−2^ s^−1^. Growth conditions used in different experiments are detailed in sections below.

F2 *DEK1* modulated lines crossed with GFP:CESA3 were used for CESA dynamics experiments. Non-genotyped segregating F2 plants were grown on MS 1% sucrose media for 3 days, imaged and then genotyped for identification of *DEK1* modulated lines containing *pCESA3:GFP:CESA3* constructs. After confocal imaging, *OE CALPAIN pCESA3:GFP:CESA3* F2 plants were transferred onto MS 1% sucrose media supplemented with 10 μg ml^-1^ hygromycin. Plants were grown horizontally for 10-14 days together with Col-0 and *GFP CESA3* plants as positive controls. Only imaging data for *OE CALPAIN pCESA3:GFP:CESA3* plants which survived the hygromycin selection were used for CESA dynamics analysis.

*dek1-4 pCESA3:GFP:CESA3* plants were transferred to MS 1% sucrose media supplemented with Plant Preservative Mixture (Plant Cell Technology, Washington DC, USA) diluted in ratio 1:1000 after CESA imaging. When plants recovered and developed first true leaves (7-10 days post imaging), DNA was isolated using Edwards buffer method (Edwards et al., 1991) and analysed by dCAPS (Neff et al., 1998) (Supplementary Table 2) and KASP genotyping (LGC Genomics, Berlin, Germany). Only imaging data for *dek1-4* homozygous plants was used in further CESA dynamics analysis.

### Analysis of *CESA* transcript levels

RNA was isolated from 6-day old *Arabidopsis* plants using the QIAGEN RNeasy Mini Kit (ID: 74104, QIAGEN, Hilden, Germany). RNA quality was determined using a Nanodrop 2000/2000c spectrophotometer (Thermo Fisher Scientific, Waltham, MA, USA). For relative quantification (RT)-qPCR experiments, cDNA synthesis was performed using Maxima First Strand cDNA Synthesis Kit (K1641, Thermo Fisher Scientific, Waltham, MA, USA) using 1 μg of RNA. Power SYBR Green Mastermix (4367659, Thermo Fisher Scientific, Waltham, MA, USA) was used for RT-qPCR and runs performed in 384 well plates (AB3384, Thermo Fisher Scientific, Waltham, MA, USA) in 5 μl reactions. Primers used for RT-qPCR are outlined in Supplementary Table 2. Lightcycler ABI 7900 HT (Applied Biosystems, Foster City, California, United States) lightcycler was used for RT-qPCR using amplification, conditions: 50° C – 2 min; 95° C – 10 min; 40 cycles of 95° C for 15 s, 60° C for 1 min; 95° C for 15 s, 60° C for 15 s, 95° for 15 s. qPCR data was analyzed using SDS v 2.4.1 analysis software (Thermo Fisher Scientific, Waltham, MA, USA). *CESA* expression levels were normalized against *GLYCERALDEHYDE-3-PHOSPHATE DEHYDROGENASE* (*GADPH*) which was used as a reference gene.

Relative gene expression values expressed as fold changes were calculated the method of (Livak and Schmittgen, 2001). *CESA* expression levels were treated as 1 in Col-0 WT and values in *DEK1* modulated lines were calculated compared to Col-0. Gene expression fold changes lower than 0.5 and higher than 2.5, compared to Col-0, were treated as significant.

### Cell wall composition analysis

Cell wall composition analysis was performed on pooled 10-day old whole seedlings. Analysis of the cell wall polysaccharide composition of alcohol insoluble cell wall residues (AIR) Col-0 and *DEK1* modulated lines was performed according to (Pettolino et al., 2012). Analysis of the total crystalline cellulose in seedlings was performed using Updegraff assay (Updegraff, 1969) of AIR (10-50 mg).

### Immunofluorescence labelling

Cotyledon tissues from 10 day old seedlings were fixed and embedded according to (Wilson and Bacic, 2012). A Leica Ultracut R microtome (Leica Microsystems, Germany) was used to obtain 250 nm thin sections and labelled with JIM5 (low methyl esterified HG; (Knox et al., 1990)), JIM7 (high methyl esterified HG; (Knox et al., 1990)), and LM15 (xyloglucan; (Marcus et al., 2008)) antibodies at 1:10 dilution, secondary antibody was Alexa Fluor 488 goat anti-rat IgG (H+L) (Life Technology; # A48262) with 1:100 dilutions. For crystalline cellulose labelling, CBM3a (Blake et al., 2006) was used at 1:50 dilution, the secondary antibody used was anti-6x-His tag monoclonal (Invitrogen, # MA1-21315) with 1:100 dilutions, the third antibody was Alexa Fluor 488 goat anti-mouse IgG (H+L) (Life Technology; # A11006) with 1:100 dilutions. Images were acquired with an Olympus BX53 microscope under GFP channel. Calcofluor white (0.02%) was used to stain cellulose and images were acquired under UV channel. Two biological replicates from two independent lines were performed.

### Transmission-electron microscopy (TEM) and immunolabelling

Thin sections (~80 nm) from cotyledon tissues were acquired as stated above. Antibody labelling and post-staining were performed according to (Wilson and Bacic, 2012). For pectin labelling, JIM5 and JIM7 antibodies were used at 1:15 dilution, secondary antibodies used was goat anti-rat 18 nm gold conjugated secondary antibody (Jackson Immuno Research #112-215-167) at 1:20 dilutions. For crystalline cellulose labelling, CBM3a at 1:50 dilution, secondary anti-6x-His tag monoclonal antibody (Invitrogen, # MA1-21315) at 1:175 dilutions, and the third antibody was goat anti-mouse 12 nm gold conjugated secondary antibody (Jackson Immuno Research #115-205-166) at 1:35 dilution. Grids were post-stained using 2% uranyl acetate for 10 min and Reynold’s lead citrate for 1 min. Grids were imaged using a Jeol 2100 EM equipped with a Gatan Orius SC 200 CCD camera. Two biological replicates from two independent lines were performed.

### Spinning disc confocal microscopy of CSC

Three-day old seedlings were mounted on microscope object slides and imaged using instruments and settings as described in (Sampathkumar et al., 2013). Photobleaching used to access CSC trafficking dynamics was performed using a FRAP/PA system (Roper Scientific, Acton, MA, USA) integrated into the spining disc confocal imaging system, as described in (Sampathkumar et al., 2013).

### CSC timelapse processing

Image processing was performed using ImageJ software (Rasband, W., NIH, Bethesda, MD, USA). Brightness and contrast of the raw time-lapses were modified. Subtract background function (rolling ball size 30-40 pixels) and Walking average plugin (three frames averaged) were used to remove background noise and to obtain clearer images.

Density of CSC particles was determined from confocal time-lapse images using IMARIS 7.4 image analysis software (Bitplane, Oxford Instruments, Oxford, United Kingdom). Time-lapse image stacks were pre-processed in ImageJ as described above and imported into IMARIS. Spot detection and tracking function of the IMARIS was used to detect and label CSC particles. CSC particle numbers used to calculate mean CSC density were taken from three random timeframes from each analyzed pavement cell form every imaged cotyledon for each of the analyzed genotypes. Density was calculated as mean of CSC particles density per one μm^2^ for each cell, plant and genotype.

Velocity analysis of CESA particles imaged using confocal microscopy was performed using Fluorescent Image Evaluation Software for Tracking and Analysis (FIESTA) software (Ruhnow et al., 2011), which is available under open license from FIESTA, University of Dresden Fusion Forge Wiki webpage (https://fusionforge.zih.tu-dresden.de/plugins/mediawiki/wiki/fiesta/index.php/FIESTA). Raw confocal time-lapse images were imported into FIESTA. Maximum signal intensity projections were used to generate kymographs. Kymographs were generated from the CSC trajectories manually drawn using segmented line tool in FIESTA.

Analysis of the data obtained by FRAP was performed using a modified version of the CSC density measurement method described above. CSC particles detection and quantification was performed on 9 x 9 μm cell patches rather than 10 x 10 μm, which were the dimensions of bleached surface. After performing detection of fluorescently labelled CSC particles using IMARIS software, CSC post-FRAP density, recovery rates and average plasma membrane life-time of the complexes was calculated as described in (Sampathkumar et al., 2013).

### Cell wall isolation for AFM topographical imaging and nanoindentation

Cotyledon pavement epidermal cell walls were isolated for analysis with the atomic force microscope (imaging and nanoindentation). Rectangular microscope coverslips (Roth, Karlsruhe, Germany) were covered with 200 μl of 0.01% poly-D-lysine-hydrobromide (PDL; P6407, Sigma Aldrich, St. Luis, MO, USA) in deionised water. Coverslips were left for 15-20 min to allow the PDL to polymerize, after which coverslips were washed thoroughly with deionized water and dried in a stream of nitrogen.

We used 10-day old seedlings of *Arabidopsis* to obtain cotyledons for cell wall isolation. The larger of the two cotyledons was cut off using surgical tweezers, placed on a PDL coated coverslip (with the abaxial side facing the coverslip) and the following steps were performed under a stereomicroscope (Zeiss Stemi 508, Zeiss, Jena, Germany). The cotyledon was pressed gently on one side using surgical tweezers and then a vertical cut was made using a micro-surgery scalpel (Fine Scientific Tools, Vancouver, Canada), across the middle of the cotyledon in a direction perpendicular to its longer axis. After the vertical cut, a second cut was performed in parallel to the longer cotyledon axis to remove all leaf tissues and expose the abaxial epidermis with patches of the cell wall monolayers on its edge (Supplementary Figure 6). Regions with cell wall monolayers were observable under 50x magnification as small opaque patches of material on the edge of the abaxial epidermis (Supplementary Figure 6).

The remaining parts of the dissected cotyledon were carefully attached to the surface with nitrocellulose based red stained adhesive (nail polish) to immobilize the sample during imaging and mechanical probing. Adhesive was applied with an injection syringe needle, carefully avoiding regions with cell wall monolayers and left to cure for 5-10 seconds. Samples were then incubated with 500 μl of 1% SDS for 20-30 s to remove organelles, proteins and other cellular components released during the cotyledon dissection. SDS was removed by washing with deionized water, ensuring that the dissected cotyledon does not curl up on the coverslip. For high-resolution AFM topological imaging the samples were left overnight to completely dry while they were strored in water for mechanical measurements to prevent loss of native mechanicl properties.

### AFM topographical imaging

For measuring the CMF diameter, a Dimension 3100 AFM (Veeco, Plainview, NY, USA) was used. Imaging was performed in tapping mode in air, using silicon ARROW-NCR cantilevers (tip radius <10 nm; Nanoworld, Neuchâtel, Switzerland).The optical camera integrated into the AFM setup was used to identify regions of interest for imaging. Regions with a size of 2 x 2 μm^2^ were imaged with a resolution of 512 x 512 pixels. We recorded height channel images (Supplementary Figure 8) and amplitude channel images (Figure 4A-C). Imaging of all genotypes was performed in biological duplicate; that is, the same experiments were repeated twice with two sets of plants grown independently at different times.

### Analysis of the cellulose microfibril images

High-resolution, high-quality cell wall monolayers imaged in dry conditions were analyzed using custom made MATLAB script to determine the diameter of the imaged CMFs. For the analysis of CMF diameter, we used the height channel AFM images (Supplementary Figure 8) as obtained from the AFM software. The extracted values of the CMF diameter were pooled for each genotype and CMF diameter values.

### Mechanical characterization of isolated cell walls using atomic force microscopy

To determine the apparent elastic modulus of the isolated cell wall monolayers, we performed AFM-based nanoindentation (manual force mapping mode), applying a method previously described (Yakubov et al., 2016) and modified for our experimental conditions. Nanoindentation experiments were performed using a JPK Nanowizard 3 AFM (Bruker Nano GmbH, Berlin, Germany) mounted on an Olympus IX71 inverted phase-contrast microscope (Olympus Corporation, Shinjuku City, Tokyo, Japan). Coverslips with prepared samples were mounted onto the AFM sample holder as described in (Yakubov et al., 2016).

An optical CCD camera integrated into the AFM setup was used during the entire measurement; first to align the cantilever and then to perform constant checks of the sample and cantilever properties. We used DNP-10 cantilevers (cantilever A) with a nominal spring constant of 0.35 N m^-1^ (Bruker Corporation, Billerica, MA, USA). The optical sensitivity of the cantilevers was determined from contactless oscillations after which the thermal noise method was used to obtain the spring constant. The used cantilevers had spring constants ranging from 0.30-0.38 N m^-1^.

Before starting the mechanical characterization, topological imaging of the cell wall monolayer was performed to identify the thinnest regions of the sample. First, using intermittent contact mode, we imaged an area of 5 x 5 μm^2^ with a resolution of 512 x 512 pixels. Imaging of a 5 x 5 μm^2^ region was always perfomed near the edge of the dissected cotyledon, so that part of imaged area represented the bare glass slide to which the cotyledon was attached. The flat glass slide surface, which contained some tissue debris after dissection, was used as a reference (position 0 nm), to determine the thickness of the imaged sample (Supplementary Figure 7). If sample thickness was in the 0.5-2.0 μm range, we selected a smaller 1 x 1 μm^2^ region within the original 5 x 5 μm area and repeated the topography imaging to re-confirm cell wall thickness (Supplementary Figure 7). If the thickness of the topological features inside this region remained in the 0.5-2 μm range, we performed nanoindentation on that region. This range of sample thicknesses values was empirically chosen. During method development, we determined that regions thicker than 2.0 μm were either wrinkled cell wall patches or cell wall bilayers (i.e. from both adaxial and abaxial sides; thickness 2.5-4 μm). Samples thinner than 0.5 μm were not used to avoid non-specific surface interactions and to minimize the contribution of the underlying glass surface to the elastic modulus.

For nanoindentation, the 1 x 1 μm^2^ region of interest was further divided into a 16 x 16 array. Each position was indented once to obtain a force map consisting of 256 force-displacement curves. For each approach-retract cycle, the contact force was set to 2.5 nN and the z-piezo speed was 1 μm s^-1^. The average indentation depth was 180 nm for Col-0, 164 nm for *OE CALPAIN* and 147 nm for *dek1-4*, while indentation depth values where mostly in the range of 100-350 nm (Supplementary Figure 9G). While the indentation depths partially exceeded 10% of the sample thickness, the values lie in a similar range. Contributions from the underlying glass surface can thus be expected to be similar for all three samples.

One force map was obtained from each dissected cotyledon. Imaging of all genotypes was performed in biological duplicate; the same experiments were repeated two times, with two sets of plants grown independently at different times. From each replicate, cotyledons from 4-6 different plants were imaged, one cotyledon per plant. Force maps were analyzed using JPK SPM processing software, version 6.1.9 (Bruker Nano GmbH). As adhesion was observed in the retract segments of the force-displacement curves, the approach segments were fit with the Hertz model to obtain the apparent elastic modulus. The Poisson ratio was set to 0.5 and the baseline value was unpinned and set to 0. The extracted apparent elastic moduli from different cotyledons for the same genotype were pooled and *dek1-4* and *OE CALPAIN* values were compared with Col-0.

### Cell wall integrity assays

Plants used for the Isoxaben treatments were sterilized and grown for 6 days on MS media containing 1% sucrose (as described in Section 2.3), with the addition of 2 nM Isoxaben (Sigma-Aldrich, St. Louis, MO, USA) dissolved in DMSO (Honeywell, Charlotte, NC, USA), and on MS 1% sucrose containing media supplemented with DMSO as a control. Experiments were performed in biological duplicates and technical triplicates. Salt stress experiments were performed as described in (Feng et al., 2018). Plants for osmotic treatments were grown for 6 days on MS 1% sucrose media supplemented with 200 mM mannitol or control plates. Each line and treatment were grown in biological duplicate and technical triplicate. Ectopic lignification experiments were performed as described in (Chaudhary et al., 2020).

Plants were imaged using a Keyence VHX-6000 digital microscope (Keyence Corporation of America, Itasca, IL, USA). Whole plates were placed on a motorized microscope stage and series of images were acquired in sequence covering the entire surface of the plate, and then automatically stitched by the software to generate a single whole-plate image. We used objective ZS20, 20X magnification, on black field, with epi-illumination, for imaging 6-day Isoxaben grown plants, and NaCl and mannitol treated plants. For plants grown overnight on Isoxaben for lignin staining, we used ZS20 objective, 200X magnification, on white field without epi-illumination. Root and hypocotyl lengths were quantified using Measure length function of ImageJ software (Rasband, W., NIH, Bethesda, MD, USA).

### Statistical analysis

Unless stated otherwise, all statistical analysis in this study, except the data from cell wall integrity assays, were performed using unpaired Student t-test, results were treated statistically significant if *p*<0.05. We used online Student t-test calculator at the webpage https://www.graphpad.com/quickcalcs/ttest1.cfm.

Analysis of data from cell wall integrity assyas - Isoxaben, NaCl, Mannitol growth assays and lignin staining was performed using estimation statistics. After obtaining root length data and quantifying signal intensity of lignin staining using ImageJ, statistical significance of the results was assessed using the online, open-access Estimation Stats software (https://www.estimationstats.com/#/) and data are plotted using Cumming estimation plot as described in (Ho et al., 2019).

## AUTHOR CONTRIBUTIONS

LN performed CSC imaging, AFM measurements and imaging and CWI sensing experiments, and analysed all the data from these experiments. KLJ, TB and AS conceptualized and designed the research. LN, GEY, KLJ and KGB conceptualized, developed, and validated cell wall isolation method and AFM imaging and force spectroscopy protocols. YM performed immunofluorescences and immunohistochemical imaging experiments and analysed the data. GEY wrote the MATLAB script for analysis of AFM cellulose images and made *DEK1* crosses with CSC reporter line. AS crossed *DEK1* modulated lines into CSC reporter line. LN wrote the original draft of the article manuscript. KLJ, TB, KGB and AS edited, curated and corrected the original draft of the manuscript. KB, GEY, and YM gave valuable input and comments on the article manuscript.

## ACKNOWLEDGMENTS

LN was funded by University of Melbourne Research Scholarship, Max Planck Society, Norma Hilda Schuster neé Swift Scholarship and Albert Shimmins Postgraduate Writing-up Award. YM acknowledges the support of a University of Melbourne Research Scholarship. KLJ and AB would like to acknowledge the support of funds from the ARC Centre of Excellence in Plant Cell Walls (CE1101007) and start-up funds from La Trobe University. Authors would like to thank and acknowledge Mr Pengfei (Alfie) Hao of La Trobe University, Australia, for performing cell wall composition and data analysis and the La Trobe BioImaging facility for assistance with electron microscopy. Authors would like to acknowledge A/Prof Monika S Doblin of La Trobe University, Australia for critical discussion and advice on method development and data interpretation. Finally, authors would like to acknowledge Ms. Reinhild Dünnebacke and Ms. Irina Berndt of the Max Planck Institute of Colloids and Interfaces, Germany, for their invaluable support and assistance during the AFM imaging and mechanical characterization of the isolated cell walls.

## REFERENCES

Amanda D, Doblin MS, Galletti R, Bacic A, Ingram GC, L. JK (2016) DEFECTIVE KERNEL1 (DEK1) regulates cell walls in the leaf epidermis. Plant Physiology 172: 2204–2218

Amanda D, Doblin MS, MacMillan CP, Galletti R, Golz JF, Bacic A, Ingram GC, Johnson KL (2017) Arabidopsis DEFECTIVE KERNEL1 regulates cell wall composition and axial growth in the inflorescence stem. Plant Direct 1: e00027

Basu D, Tian L, Debrosse T, Poirier E, Emch K, Herock H, Travers A, Showalter AM (2016) Glycosylation of a Fasciclin-Like Arabinogalactan-Protein (SOS5) Mediates Root Growth and Seed Mucilage Adherence via a Cell Wall Receptor-Like Kinase (FEI1/FEI2) Pathway in Arabidopsis. PloS one 11: e0145092–e0145092

Becraft PW, Li K, Dey N, Asuncion-Crabb Y (2002) The maize dek1 gene functions in embryonic pattern formation and cell fate specification. Development 129: 5217–5225

Bischoff V, Desprez T, Mouille G, Vernhettes S, Gonneau M, Höfte H (2011) Phytochrome regulation of cellulose synthesis in Arabidopsis. Current Biology 21: 1822–1827

Blackburn MR, Haruta M, Moura DS (2020) Twenty Years of Progress in Physiological and Biochemical Investigation of RALF Peptides1 Plant Physiology 182: 1657–1666

Blake AW, McCartney L, Flint JE, Bolam DN, Boraston AB, Gilbert HJ, Knox JP (2006) Understanding the biological rationale for the diversity of cellulose-directed carbohydrate-binding modules in prokaryotic enzymes. Journal of Biological Chemistry 281: 29321–29329

Bouton S, Leboeuf E, Mouille G, Leydecker MT, Talbotec J, Granier F, Lahaye M, Höfte H, Truong HN (2002) QUASIMODO1 encodes a putative membrane-bound glycosyltransferase required for normal pectin synthesis and cell adhesion in Arabidopsis. The Plant Cell 14: 2577–2590

Bringmann M, Li E, Sampathkumar A, Kocabek T, Hauser M-T, Persson S (2012) POM-POM2/CELLULOSE SYNTHASE INTERACTING1 Is Essential for the Functional Association of Cellulose Synthase and Microtubules in Arabidopsis. The Plant Cell 24: 163–177

Chaudhary A, Chen X, Gao J, Leśniewska B, Hammerl R, Dawid C, Schneitz K (2020) The Arabidopsis receptor kinase STRUBBELIG regulates the response to cellulose deficiency. PLOS Genetics 16

Cosgrove DJ (2005) Growth of the plant cell wall. Nature Reviews Molecular Cell Biology 6: 850–861

Czechowski T, Stitt M, Altmann T, Udvardi MK, Scheible W-R (2005) Genome-Wide Identification and Testing of Superior Reference Genes for Transcript Normalization in Arabidopsis. Plant Physiology 139: 5–7

Demko V, Perroud P-F, Johansen W, Delwiche CF, Cooper ED, Remme P, Ako AE, Kugler KG, Mayer KFX, Quatrano R, Olsen O-A (2014) Genetic Analysis of DEFECTIVE KERNEL1 Loop Function in Three-Dimensional Body Patterning in <em>Physcomitrella patens. Plant Physiology 166: 903–919

Denness L, McKenna JF, Segonzac C, Wormit A, Madhou P, Bennett M, Mansfield J, Zipfel C, Hamann T (2011) Cell Wall Damage-Induced Lignin Biosynthesis Is Regulated by a Reactive Oxygen Species- and Jasmonic Acid-Dependent Process in Arabidopsis Plant Physiology 156: 1364–1374

Desprez T, Juraniec M, Crowell EF, Jouy H, Pochylova Z, Parcy F, Höfte H, Gonneau M, Vernhettes S (2007) Organization of cellulose synthase complexes involved in primary cell wall synthesis in Arabidopsis thaliana. Proceedings of the National Academy of Sciences of the United States of America 104: 15572–15577

Diotallevi F, Mulder B (2007) The cellulose synthase complex: a polymerization driven supramolecular motor. Biophysical Journal 92: 2666–2673

Du J, Kirui A, Huang S, Wang L, Barnes WJ, Kiemle SN, Zheng Y, Rui Y, Ruan M, Qi S, Kim SH, Wang T, Cosgrove DJ, Anderson CT, Xiao C (2020) Mutations in the Pectin Methyltransferase QUASIMODO2 Influence Cellulose Biosynthesis and Wall Integrity in Arabidopsis. The Plant Cell 32: 3576–3597

Edwards K, Johnstone C, Thompson C (1991) A simple and rapid method for the preparation of plant genomic DNA for PCR analysis. Nucleic Acids Research 19: 1349

Engelsdorf T, Hamann T (2014) An update on receptor-like kinase involvement in the maintenance of plant cell wall integrity. Annals of Botany 114: 1339–1347

Fagard M, Desnos T, Desprez T, Goubet F, Refregier G, Mouille G, McCann M, Rayon C, Vernhettes S, Höfte H (2000) PROCUSTE1 encodes a cellulose synthase required for normal cell elongation specifically in roots and dark-grown hypocotyls of Arabidopsis. The Plant Cell 12: 2409–2424

Feng W, Kita D, Peaucelle A, Cartwright HN, Doan V, Duan Q, Liu MC, Maman J, Steinhorst L, Schmitz-Thom I, Yvon R, Kudla J, Wu H-M, Cheung AY, Dinneny JR (2018) The FERONIA receptor kinase maintains cell-wall integrity during salt stress through Ca signaling. Current Biology 28: 1–10

Fujita M, Himmelspach R, Hocart CH, Williamson RE, Mansfield SD, Wasteneys GO (2011) Cortical microtubules optimize cell-wall crystallinity to drive unidirectional growth in Arabidopsis. The Plant Journal 66: 915–928

Galletti R, Johnson KL, Scofield S, San-Bento R, Watt AM, Murray JAH, Ingram GC (2015) DEFECTIVE KERNEL 1 promotes and maintains plant epidermal differentiation. Development 142: 1978–1983

Galletti R, Verger S, Hamant O, Ingram GC (2016) Developing a ‘thick skin’: a paradoxical role for mechanical tension in maintaining epidermal integrity? Development 143: 3249–3258

García Díaz BE, Gauthier S, Davies PL (2006) Ca2+ Dependency of Calpain 3 (p94) Activation. Biochemistry 45: 3714–3722

Gjetting SK, Mahmood K, Shabala L, Kristensen A, Shabala S, Palmgren M, Fuglsang AT (2020) Evidence for multiple receptors mediating RALF-triggered Ca2+ signaling and proton pump inhibition. The Plant Journal 104: 433–446

Haas KT, Wightman R, Meyerowitz EM, Peaucelle A (2020) Pectin homogalacturonan nanofilament expansion drives morphogenesis in plant epidermal cells. Science 367: 1003–1007

Hamann T (2014) The Plant Cell Wall Integrity Maintenance Mechanism—Concepts for Organization and Mode of Action. Plant and Cell Physiology 56: 215–223

Hamant O, Heisler MG, Jönsson H, Krupinski P, Uyttewaal M, Bokov P, Corson F, Sahlin P, Boudaoud A, Meyerowitz EM, Couder Y, Traas J (2008) Developmental patterning by mechanical signals in Arabidopsis. Science 322: 1650–1655

Heim DR, Skomp JR, Tschabold EE, Larrinua IM (1990) Isoxaben Inhibits the Synthesis of Acid Insoluble Cell Wall Materials In Arabidopsis thaliana. Plant Physiology 93: 695–700

His I, Driouich A, Nicol F, Jauneau A, Höfte H (2001) Altered pectin composition in primary cell walls of korrigan, a dwarf mutant of Arabidopsis deficient in a membrane-bound endo-1,4-beta-glucanase. Planta 212: 348–358

Ho J, Tumkaya T, Aryal S, Choi H, Claridge-Chang A (2019) Moving beyond P values: data analysis with estimation graphics. Nature Methods 16: 565–566

Hu H, Zhang R, Tang Y, Peng C, Wu L, Feng S, Chen P, Wang Y, Du X, Peng L (2019) Cotton CSLD3 restores cell elongation and cell wall integrity mainly by enhancing primary cellulose production in the Arabidopsis cesa6 mutant. Plant Molecular Biology 101: 389–401

Johnson KL, Degnan KA, Ross Walker J, Ingram GC (2005) AtDEK1 is essential for specification of embryonic epidermal cell fate. The Plant Journal 44: 114–127

Johnson KL, Faulkner C, Jeffree CE, Ingram GC (2008) The phytocalpain defective kernel 1 is a novel Arabidopsis growth regulator whose activity is regulated by proteolytic processing. The Plant Cell 20: 2619–2630

Johnson KL, Faulkner, C., Jeffree, C. E., Ingram, G. C. (2008) The phytocalpain defective kernel 1 is a novel Arabidopsis growth regulator whose activity is regulated by proteolytic processing. The Plant Cell 20: 2619–2630

Johnson KL, Gidley MJ, Bacic A, Doblin MS (2018) Cell wall biomechanics: a tractable challenge in manipulating plant cell walls ‘fit for purpose’! Current Opinion in Biotechnology 49: 163–171

Kafle K, Xi X, Lee CM, Tittmann BR, Cosgrove DJ, Park YB, Kim SH (2014) Cellulose microfibril orientation in onion (Allium cepa L.) epidermis studied by atomic force microscopy (AFM) and vibrational sum frequency generation (SFG) spectroscopy. Cellulose 21: 1075–1086

Knox JP, Linstead PJ, King J, Cooper C, Roberts K (1990) Pectin esterification is spatially regulated both within cell walls and between developing tissues of root apices. Planta 181: 512–521

Kohorn BD, Johansen, S., Shishido, A., Todorova, T., Martinez, R., Defeo, E., Obregon, P. (2009) Pectin activation of MAP kinase and gene expression is WAK2 dependent. The Plant Journal 60: 974–982

Lane DR, Wiedemeier A, Peng L, Höfte H, Vernhettes S, Desprez T, Hocart CH, Birch RJ, Baskin TI, Burn JE, Arioli T, Betzner AS, Williamson RE (2001) Temperature-sensitive alleles of RSW2 link the KORRIGAN endo-1,4-beta-glucanase to cellulose synthesis and cytokinesis in Arabidopsis. Plant Physiol 126: 278–288

Lid SE, Gruis D, Jung R, Lorentzen JA, Ananiev E, Chamberlin M, Niu X, Meeley R, Nichols S, Olsen O-A (2002) The defective kernel 1 (dek1) gene required for aleurone cell development in the endosperm of maize grains encodes a membrane protein of the calpain gene superfamily. Proceedings of the National Academy of Sciences of the United States of America 99: 5460–5465

Livak KJ, Schmittgen TD (2001) Analysis of relative gene expression data using real-time quantitative PCR and the 2^(-ΔΔCt)^ Method. Methods 25

Lopez-Sanchez P, Cersosimo J, Wang D, Flanagan B, Stokes JR, Gidley MJ (2015) Poroelastic Mechanical Effects of Hemicelluloses on Cellulosic Hydrogels under Compression. PLOS one 10:e0122132

Marcus SE, Verhertbruggen Y, Herve C, Ordaz-Ortiz JJ, Farkas V, Pedersen HL, Willats WG, Knox JP (2008) Pectic homogalacturonan masks abundant sets of xyloglucan epitopes in plant cell walls. BMC Plant Biology 8: 60

Mouille G, Ralet MC, Cavelier C, Eland C, Effroy D, Hématy K, McCartney L, Truong HN, Gaudon V, Thibault JF, Marchant A, Höfte H (2007) Homogalacturonan synthesis in Arabidopsis thaliana requires a Golgi-localized protein with a putative methyltransferase domain. The Plant Journal 50: 605–614

Moulia B, Douady S, Hamant O (2021) Fluctuations shape plants through proprioception. Science 372:eabc6868

Neff MM, Neff JD, Chory J, Pepper AE (1998) dCAPS, a simple technique for the genetic analysis of single nucleotide polymorphisms: experimental applications in Arabidopsis thaliana genetics. The Plant Journal 14: 387–392

Nixon BT, Mansouri K, Singh A, Du J, Davis JK, Lee J-G, Slabaugh E, Vandavasi VG, O’Neill H, Roberts EM, Roberts AW, Yingling YG, Haigler CH (2016) Comparative Structural and Computational Analysis Supports Eighteen Cellulose Synthases in the Plant Cellulose Synthesis Complex. Scientific Reports 6: 28696

Ono Y, Sorimachi H (2012) Calpains: an elaborate proteolytic system. Biochim Biophys Acta 1824: 224–236

Paredez AR, Somerville CR, Ehrhardt DW (2006) Visualization of cellulose synthase demonstrates functional association with microtubules. Science 312: 1491–1495

Persson Sea (2007) Genetic evidence for three unique components in primary cell-wall cellulose synthase complexes in Arabidopsis. Proc. Natl. Acad. Sci. U.S.A. 104: 15566–15571

Pettolino FA, Walsh C, Fincher GB, Bacic A (2012) Determining the polysaccharide composition of plant cell walls. Nat Protoc 7: 1590–1607

Polko JK, Kieber JJ (2019) The Regulation of Cellulose Biosynthesis in Plants. The Plant Cell 31: 282–296

Roeder AHK, Cunha A, Ohno CK, Meyerowitz EM (2012) Cell cycle regulates cell type in the Arabidopsis sepal. Development 139: 4416–4427

Ruhnow F, Zwicker D, Diez S (2011) Tracking single particles and elongated filaments with nanometer precision. Biophysical Journal 100: 2820–2828

Sampathkumar A, Gutierrez R, McFarlane HE, Bringmann M, Lindeboom J, Emons A-M, Samuels L, Ketelaar T, Ehrhardt DW, Persson S (2013) Patterning and Lifetime of Plasma Membrane-Localized Cellulose Synthase Is Dependent on Actin Organization in Arabidopsis Interphase Cells. Plant Physiology 162: 675–688

Sánchez-Rodríguez C, Ketelaar K, Schneider R, Villalobos JA, Somerville CR, Persson S, Wallace IS (2017) BRASSINOSTEROID INSENSITIVE2 negatively regulates cellulose synthesis in Arabidopsis by phosphorylating cellulose synthase 1. Proc. Natl. Acad. Sci. U.S.A. 114: 3533–3538

Shim I, Law R, Kileeg Z, Stronghill P, Northey JGB, Strap JL, Bonetta DT (2018) Alleles Causing Resistance to Isoxaben and Flupoxam Highlight the Significance of Transmembrane Domains for CESA Protein Function. Frontiers in Plant Science 9

Stegmann M, Monaghan J, Smakowska-Luzan E, Rovenich H, Lehner A, Holton N, Belkhadir Y, Zipfel C (2017) The receptor kinase FER is a RALF-regulated scaffold controlling plant immune signaling. Science 355: 287–289

Tenhaken R (2014) Cell wall remodeling under abiotic stress. Frontiers in Plant Science 5: 771

Tonami K, Kurihara Y, Arima S, Nishiyama K, Uchijima Y, Asano T, Sorimachi H, Kurihara H (2011) Calpain-6, a microtubule-stabilizing protein, regulates Rac1 activity and cell motility through interaction with GEF-H1. Journal of Cell Science 124: 1214–1223

Tran D, Galletti R, Neumann ED, Dubois A, Sharif-Naeini R, Geitmann A, Frachisse JM, Hamant O, Ingram GC (2017) A mechanosensitive Ca2+ channel activity is dependent on the developmental regulator DEK1. Nature Communications 8

Updegraff DM (1969) Semimicro determination of cellulose in biological materials. Analytical Biochemistry 32: 420–424

Vaahtera L, Schulz J, Hamann T (2019) Cell wall integrity maintenance during plant development and interaction with the environment. Nature Plants 5: 924–932

Vain T, Crowell EF, Timpano H, Biot E, Desprez T, Mansoori N, Trindade LM, Pagant S, Robert S, Höfte H, Gonneau M, Vernhettes S (2014) The Cellulase KORRIGAN Is Part of the Cellulose Synthase Complex. Plant Physiology 165: 1521–1532

Verger S, Long Y, Boudaoud A, Hamant O (2018) A tension-adhesion feedback loop in plant epidermis. eLife 7: e34460

Wang C, Barry JK, Min Z, Tordsen G, Rao AG, Olsen OA (2003) The calpain domain of the maize DEK1 protein contains the conserved catalytic triad and functions as a cysteine proteinase. Journal of Biological Chemistry 278: 34467–34474

Wang T, Park YB, Cosgrove DJ, Hong M (2015) Cellulose-Pectin Spatial Contacts Are Inherent to Never-Dried Arabidopsis Primary Cell Walls: Evidence from Solid-State Nuclear Magnetic Resonance Plant Physiology 168: 871–884

Wang X, Wilson L, Cosgrove DJ (2020) Pectin methylesterase selectively softens the onion epidermal wall yet reduces acid-induced creep. Journal of Experimental Botany 71: 2629–2640

Wilson SM, Bacic A (2012) Preparation of plant cells for transmission electron microscopy to optimize immunogold labeling of carbohydrate and protein epitopes. Nature Protocols 7: 1716–1727

Xiao C, Anderson CT (2016) Interconnections between cell wall polymers, wall mechanics, and cortical microtubules: Teasing out causes and consequences. Plant signaling & behavior 11: e1215396–e1215396

Xiao C, Zhang T, Zheng Y, Cosgrove DJ, Anderson CT (2016) Xyloglucan Deficiency Disrupts Microtubule Stability and Cellulose Biosynthesis in Arabidopsis, Altering Cell Growth and Morphogenesis. Plant Physiology 170: 234–249

Yakubov GE, Bonilla MR, Chen H, Doblin MS, Bacic A, Gidley MJ, Stokes JR (2016) Mapping nano-scale mechanical heterogeneity of primary plant cell walls. Journal of Experimental Botany 67: 2799–2816

Yang Y, Yu Y, Liang Y, Anderson CT, Cao J (2018) A Profusion of Molecular Scissors for Pectins: Classification, Expression, and Functions of Plant Polygalacturonases. Frontiers in Plant Science 9: 1208

Zhang T, Tang H, Vavylonis D, Cosgrove DJ (2019) Disentangling loosening from softening: insights into primary cell wall structure. The Plant Journal 100: 1101–1117

Zhang T, Zheng Y, Cosgrove DJ (2016) Spatial organization of cellulose microfibrils and matrix polysaccharides in primary plant cell walls as imaged by multichannel atomic force microscopy. The Plant Journal 85: 179–192

Zhang X, Yang Z, Wu D, Yu F (2020) RALF–FERONIA Signaling: Linking Plant Immune Response with Cell Growth. Plant Communications 1: 100084

Zhu X, Li S, Pan S, Xin X, Gu Y (2018) CSI1, PATROL1, and exocyst complex cooperate in delivery of cellulose synthase complexes to the plasma membrane. Proceedings of the National Academy of Sciences 115: E3578–E3587

